# Functional connectivity predicts changes in attention over minutes, days, and months

**DOI:** 10.1101/700476

**Authors:** Monica D. Rosenberg, Dustin Scheinost, Abigail S. Greene, Emily W. Avery, Young Hye Kwon, Emily S. Finn, Ramachandran Ramani, Maolin Qiu, R. Todd Constable, Marvin M. Chun

## Abstract

The ability to sustain attention differs across people and changes within a single person over time. Although recent work has demonstrated that patterns of functional brain connectivity predict individual differences in sustained attention, whether these same patterns capture fluctuations in attention in single individuals remains unclear. Here, across five independent studies, we demonstrate that the sustained attention connectome-based predictive model (CPM), a validated model of sustained attention function, generalizes to predict attention changes across minutes, days, weeks, and months. Furthermore, the sustained attention CPM is sensitive to within-subject state changes induced by propofol as well as sevoflurane, such that individuals show functional connectivity signatures of stronger attentional states when awake than when under deep sedation and light anesthesia. Together these results demonstrate that fluctuations in attentional state reflect variability in the same functional connectivity patterns that predict individual differences in sustained attention.

## Introduction

As anyone who has struggled to sit through an esoteric film or reached the end of a paragraph without comprehending its content recognizes, we don’t sustain a continuous level of attention at every point in time. Rather, we are frequently distracted by our external environment and our own internal thoughts, and our level of focus fluctuates—intentionally or not (Seli et al., 2016)— with factors including mindlessness, motivation, resource allocation, and arousal (Esterman and Rothlein, 2019).

Functional MRI (fMRI) studies in humans have linked these moment-to-moment attention fluctuations to ongoing activity in large-scale brain networks including the default mode, dorsal attention, and salience networks (Christoff et al., 2009; Fortenbaugh et al., 2018; Kucyi et al., 2016a; Leber, 2010; Rosenberg et al., 2015; Weissman et al., 2006). A growing body of work has also related changes in fMRI and intracranial electroencephalography functional connectivity to changes in attentional and cognitive states (Gonzalez-Castillo et al., 2015; Kucyi et al., 2016b, 2018; Shappell et al., 2019; Shine et al., 2016a; Turnbull et al., 2019). The degree to which functional connectivity dynamics reflect cognitive state dynamics rather than physiological and measurement noise, however, is still debated (Calhoun et al., 2014; Gonzalez-Castillo and Bandettini, 2018; Hutchison et al., 2013; Lurie, D. J. et al., 2018; Preti et al., 2017). Furthermore, despite these advances, cognitive neuroscience lacks a comprehensive, quantitative measure of intra-individual differences in sustained attention, or changes in attention over time.

In contrast to the discussions on dynamics, there is growing consensus that models based on individuals’ unique patterns of static functional brain connectivity can predict individual differences in abilities including fluid intelligence (Finn et al., 2015; Greene et al., 2018), working memory (Avery et al.; Galeano Weber et al., 2017; Yamashita et al., 2018), and attention (Kessler et al., 2016; O’Halloran et al., 2018; Poole et al., 2016; Rosenberg et al., 2017; Yoo et al., 2018). The most extensively validated connectome-based predictive model (CPM), the sustained attention CPM (Rosenberg et al., 2016a), has generalized across six independent data sets and participant populations to predict individuals’ overall sustained attention function from data collected during rest and five different tasks (Fountain-Zaragoza et al., 2019; Jangraw et al., 2018; Rosenberg et al., 2016a, 2016b, 2018a). Derived using a data-driven technique (Shen et al., 2017), the sustained attention CPM comprises a distributed “high-attention” network of functional connections, or edges, stronger in individuals with better sustained attention function and a “low-attention” network of edges stronger in individuals with worse sustained attention.

Do these networks, which predict a person’s overall ability to maintain focus, also reflect fluctuations in attentional state? Evidence does suggest that the sustained attention CPM is sensitive to attention improvements following a pharmacological manipulation. That is, healthy adults given a single dose of methylphenidate, a common attention deficit hyperactivity disorder treatment, show higher high-attention network strength and lower low-attention network strength than unmedicated controls (Rosenberg et al., 2016b). However, this between-subjects study did not directly test whether changes in attention network strength mirror changes in attentional state, and it remains an open question whether the same functional networks that predict differences in attention *between* people also predict differences in attention *within* a single person over time.

Here, in a series of five experiments, we ask whether the sustained attention CPM predicts a person’s attention task performance—above and beyond predicting their *overall* level of sustained attention function—during short experimental blocks and functional MRI sessions spread across days, weeks, and months. Furthermore, we evaluate the model’s sensitivity to cognitive and attentional state changes resulting from pharmacological interventions by comparing task-free functional connectivity before and after the administration of two anesthetics, propofol and sevoflurane, with different mechanisms of action (for reviews see Patel and Goa, 1996; Trapani et al., 2000). Our results replicate findings that the sustained attention CPM generalizes to novel individuals to predict their average sustained attention function, and moreover demonstrate that the model predicts minute-by-minute, day-by-day, and drug-induced changes in attentional state. Thus, the same neuromarker predicts both inter- and intra-individual differences in sustained attention, and fluctuations in functional connectivity around a person’s mean “functional connectome fingerprint” in part reflect fluctuations in behaviorally relevant attentional states.

## Results

### Experiment 1: Sustained attention network strength predicts minute-to-minute attention fluctuations

As a first step, to test whether connectome-based models predict fluctuations in sustained attention, we re-analyzed functional MRI data from 25 individuals performing a challenging sustained attention task (the gradual-onset continuous performance task, or gradCPT; Esterman et al., 2013) reported in previous work (Rosenberg et al., 2016). Each participant performed up to three runs of the gradCPT during fMRI, and each run included four 3-min task blocks separated by 32-sec rest breaks. During the task, participants saw city and mountain photographs continuously transitioning from one to the next at a rate of 800 ms/image, and were instructed to press a button in response to city scenes (90%) but not to mountain scenes (10%).

Mean gradCPT sensitivity (*d*’) was 2.11 (*s.d*. = .92). Mean standard deviation of *d*’ across task blocks was .50, and mean coefficient of variation (standard deviation divided by the mean) of *d*’ across task blocks was 32%. Individuals’ overall *d*’ scores were inversely related to their coefficients of variation (*r_s_* = −.83, *p* = 2.09×10^−6^) but not to their standard deviation of *d*’ across task blocks (*rs* = −.22, *p* = .28). Suggesting that changes in *d*’ over time were, to some degree, consistent across participants, performance fluctuations were significantly albeit weakly correlated across individuals (mean pairwise Spearman correlation between participants’ *d*’ time-series = .083, *p* = .003 based on 10,000 permutations).

#### Predictions from task-block connectivity

To predict minute-by-minute gradCPT performance from functional connectivity patterns, multiple functional connectivity matrices were generated for each individual using a 268-node whole-brain functional atlas (Shen et al., 2013): one overall task matrix from data concatenated across task runs, up to 12 task-block matrices from volumes acquired during individual 3-min task blocks, and up to 9 rest-break matrices using volumes acquired during 32-sec rest breaks. Using leave-one-subject-out cross-validation, connectome-based models were trained to predict *d*’ using *n*−1 participants’ overall task matrices, and then applied to each of the held-out individual’s task-block matrices to generate block-specific *d*’ predictions. This prediction pipeline replicated that of Rosenberg et al. (2016a), except that models were applied to the held-out participant’s task-block matrices rather than to their overall task matrix (see *Methods* for details).

At the group level, models trained on overall task matrices generalized to predict task block-specific gradCPT performance in unseen individuals (mean within-subject Spearman correlation between predicted and observed block-wise *d*’ scores = .53, *s.d*. = .55; *t*_24_ = 5.46, *p* = 1.30×10^−5^; **Figures 1, 2**). Predictions were significant at *p* < .05 in 12 of 25 participants based on permutation tests. Importantly, these models, which predict block-specific gradCPT performance, previously generalized to predict participants’ mean performance over the course of the entire scan session (Rosenberg et al., 2016a).

**Figure 1.**
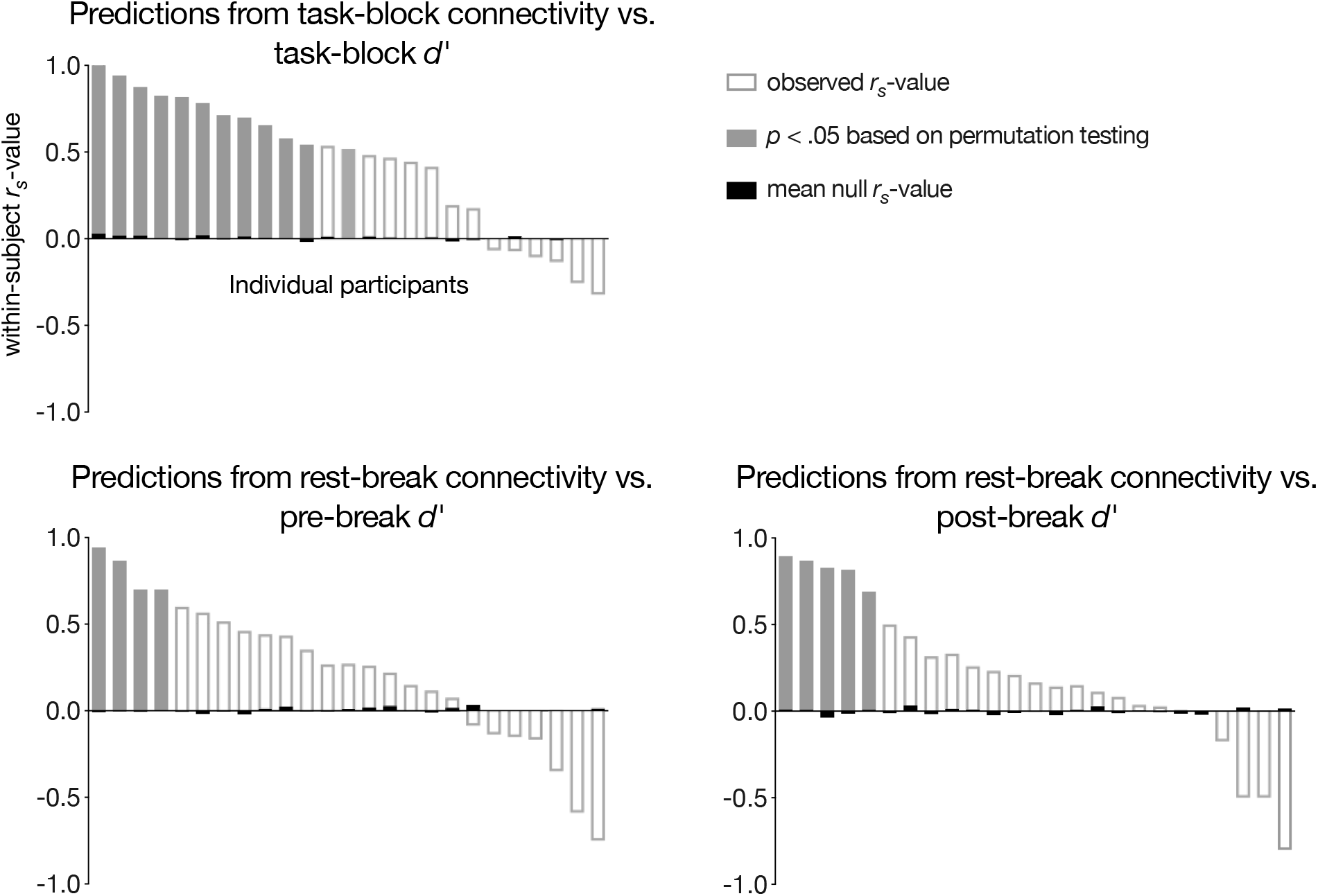
Within-subject Spearman correlations between block-wise *d*’ scores and task-block and rest-break predictions. Subject-level significance was determined with permutation testing. Group-level significance (*p* < .03 for all three models) was assessed with a *t*-test between observed and mean null correlations between predicted and observed behavior.

**Figure 2.**
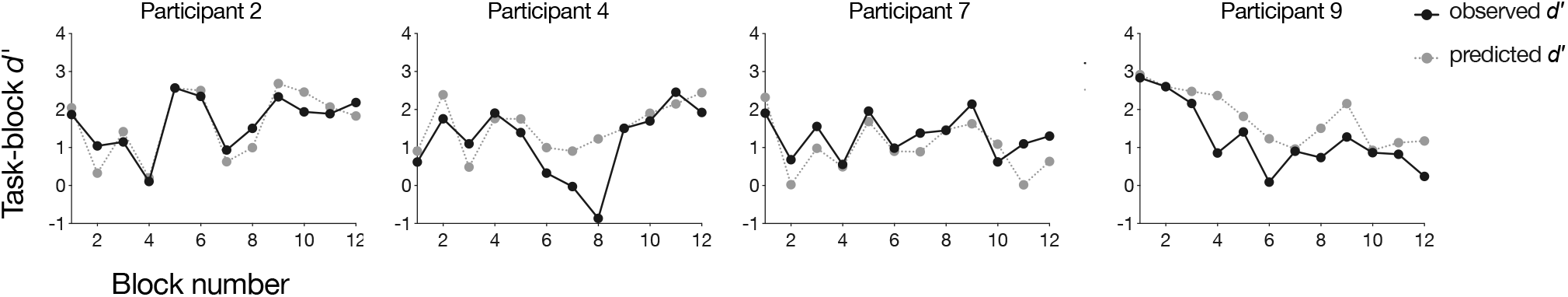
Block-wise *d*’ scores and task-block predictions from four representative individuals (participant number corresponds to the order of bars in the top panel of Figure 1).

#### Predictions from rest-break connectivity

Task-based functional connectivity measured during brief 3-min task blocks predicts block-specific task performance in individual subjects. Thus, task-free functional connectivity patterns may also reflect transient attentional states. To test this possibility, models trained on 24 participants’ overall task matrices were applied to each of the left-out subjects’ rest-break matrices to generate a predicted *d*’ score corresponding to each rest break (6 for participants with two gradCPT runs and 9 for participants with three). Because performance is not measured during rest breaks themselves, model performance was assessed by correlating predictions with the *d*’ scores from the preceding and following task blocks. These models are referred to as the “pre-break” and “post-break” models, respectively.

Models trained on overall task matrices generalized to predict left-out individuals’ performance fluctuations from functional connectivity observed during task-free rest breaks. Model predictions were significantly related to performance during the blocks immediately preceding rest breaks (mean within-subject *r_s_* = .28, *s.d*. = .52; *t*_24_ = 2.57, *p* = .017; **Figure 1**) as well as performance during the blocks immediately following breaks (mean within-subject *rs* = .26, *s.d*. = .52; *t*_24_ = 2.35, *p* = .028; **Figure 1**). Predictions were significantly related to pre-break behavior in 4 of 25 participants and to post-break behavior in 5 of 25 participants.

As expected, behavioral predictions from task-block connectivity patterns were more accurate than those from rest-break patterns (task-block vs. pre-break: *t*_24_ = 2.0, *p* = .057; task-block vs. post-break: *t*_24_ = 2.53, *p* = .018). Notably, however, predictions from rest-break connectivity were not more strongly correlated with pre-break than post-break behavior (*t*_24_ = .20, *p* = .84), suggesting that functional connectivity patterns observed during mid-task rest breaks reflect local attentional state rather than past or future attentional performance alone.

The accuracy of task-block and post-break model predictions was correlated across participants, such that individuals whose attention fluctuations were better predicted from task-block connectivity were also better predicted from rest-break connectivity (*r_s_* = .47, *p* = .018). Pre-break model predictive power was not significantly related to post-break (*r_s_* = .16, *p* = .45) or task-block model predictive power (*r_s_* = .27, *p* = .19).

#### Individual differences in model performance

Task-block models more accurately predicted attention fluctuations in participants with higher coefficients of variation (*r_s_* = .63, *p* = .001). There was a trend such that task block models also better predicted performance in individuals with lower overall *d*’ values (*r_s_* = –.46, *p* = .021), although this relationship does not survive Bonferroni correction for 6 post-hoc comparisons (3 models [task-block, pre-break, post-break] × 2 behavioral measures [*d*’, coefficient of variation]). Overall *d*’ and *d*’ coefficient of variation values were not significantly related to pre-break or post-break model accuracy (*p* values > .19).

#### Predictive network anatomy

Because feature-selection and model building steps replicated those described in Rosenberg et al. (2016a), predictive networks are identical to those reported previously. Briefly, across the 25 rounds of leave-one-out cross-validation, networks predicting better *d*’ scores included 1279–1540 functional connections (mean = 1426.7, *s.d*. = 73.9; “positive networks”), and networks predicting worse *d*’ scores included 1099–1373 functional connections (mean = 1251.1, *s.d*. = 68.1; “negative networks”). The 757 edges common to all 25 positive networks and the 630 edges common to all 25 negative networks comprise the high-attention and low-attention networks, respectively (**Figure 3**). The high- and low-attention networks are widely distributed across cortex, subcortex, and cerebellum, and are robust to computational lesioning methods that exclude predictive nodes and edges in individual brain lobes and canonical functional networks (Rosenberg et al., 2016a). Thus, the current results demonstrate that the same distributed pattern of functional brain connectivity that predicts interindividual differences in sustained attention also predicts intra-individual differences in attention.

**Figure 3.**
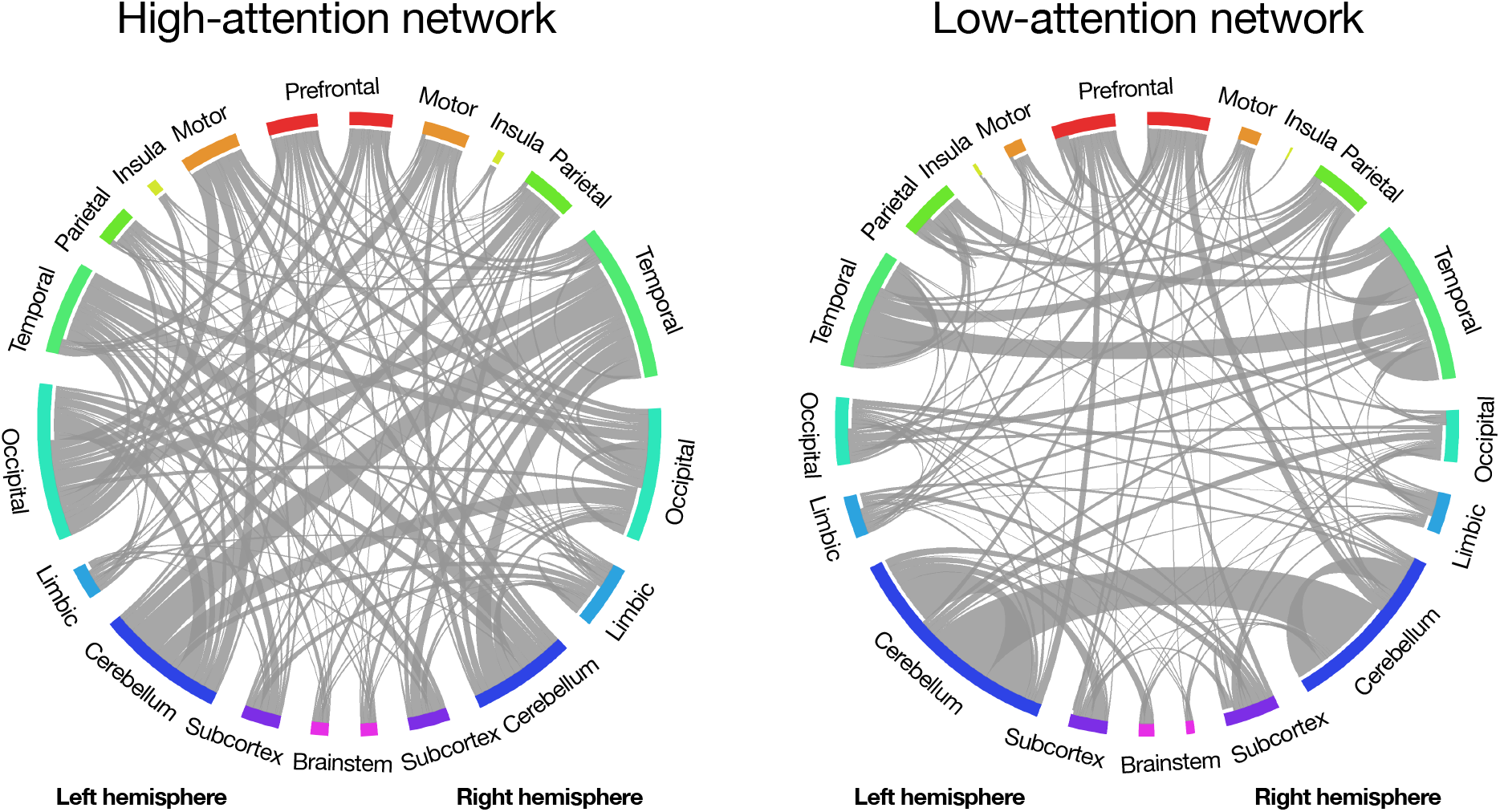
Functional connections (edges) in the high-attention and low-attention networks (Rosenberg et al., 2016a). Network nodes are grouped into macroscale brain regions; lines between them represent edges. Line width corresponds to the number of edges between region pairs. This figure was adapted with permission from Rosenberg & Chun (in press) and was created using Circos (Krzywinski et al., 2009).

### Experiment 2: Sustained attention network strength predicts session-to-session changes in attention

Experiment 1 used leave-one-subject-out cross-validation (i.e., internal validation) to demonstrate that the same functional networks that predict individual differences in sustained attention are sensitive to fluctuations in attention across 3-min task blocks. Here we test whether a model defined using the full Experiment 1 data set, the sustained attention connectome-based predictive model (CPM), predicts session-to-session variability in focus in completely new individuals. To this end, we analyzed data collected as an independent group of 50 adults performed 10 min of the gradCPT during two MRI sessions approximately three weeks apart. In this external validation sample, mean *d*’ was 2.28 (*s.d*. = .77) in session 1 and 2.14 (*s.d*. = .99) in session 2. Performance scores were correlated across sessions (*r_s_* = .72, *p* = 3.78×10^−9^) and did not significantly differ between session 1 and session 2 (*t*_49_ = 1.46, *p* = .15).

#### Attention predictions

The sustained attention CPM was applied to each participant’s session 1 and session 2 gradCPT matrices separately to predict session-specific *d*’ scores. Briefly, sustained attention CPM predictions are generated by inputting the difference between an individual’s high-attention network strength and low-attention network strength into a linear model whose coefficients were defined in previous work (Rosenberg et al., 2016a). Model outputs correspond to predicted gradCPT *d*’ scores. In the current sample, predicted *d*’ scores were significantly correlated with true *d*’ scores during the first (*r_s_* = .41, *p* = .0038) and second (*r_s_* = .70, *p* = 2.00×10^−8^) imaging sessions, demonstrating robust cross-data-set generalization (**Figure 4A**). Furthermore, predicted *d*’ was higher for participants’ better vs. worse scan session (*t*49 = 3.64, *p* = 6.56×10^−4^), and the session with the higher predicted *d*’ corresponded to the session with the higher observed *d*’ in 34 out of 50 individuals (68%; **Figure 4B**). The difference in *d*’ between participants’ first and second gradCPT sessions was also correlated with the difference in predicted *d*’ for these sessions (*r_s_* = .62, *p* = 2.26×10^−6^). Thus, the same functional connectivity patterns that predict individual differences in sustained attention reflect subtle within-subject changes in attentional performance, even in a highly reliable task.

**Figure 4.**
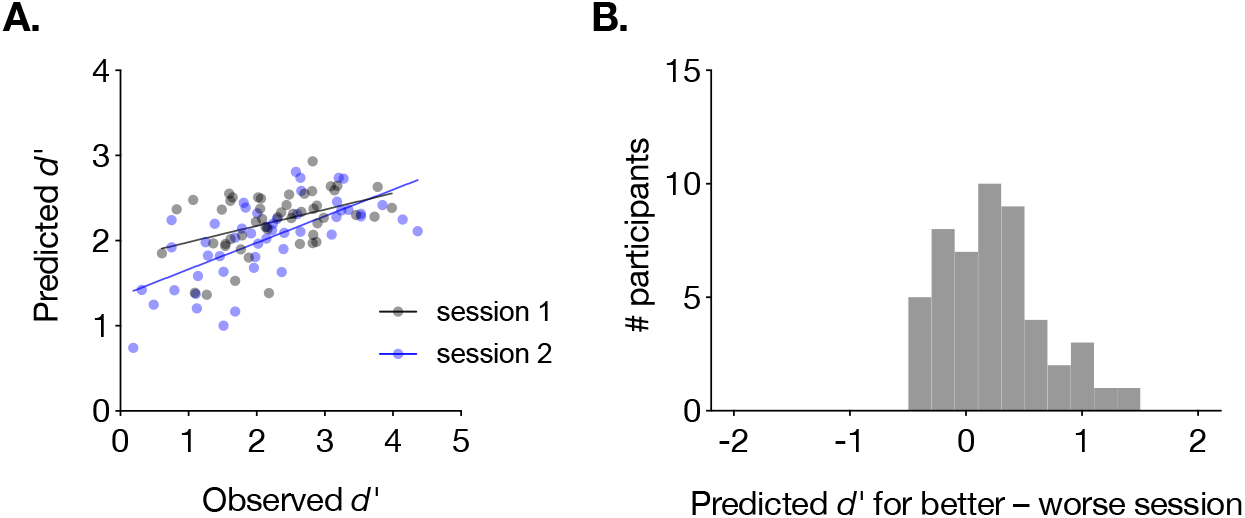
(A) The sustained attention CPM generalized to predict gradCPT performance during two neuroimaging sessions. Individual participants are represented with one gray dot corresponding to session 1 performance, and one blue dot corresponding to session 2 performance. (B) Histogram of the difference in predicted performance for each person’s better and worse task sessions. Values greater than zero indicate that the model correctly predicted a higher score for the better session.

### Experiment 3: Sustained attention network strength predicts week-to-week changes in attention in a single individual

We next investigated whether the sustained attention CPM is sensitive to within-subject variability in sustained attention over the course of weeks and months. To this end, we analyzed a longitudinal data set described in previous work (Salehi et al., 2018). This data set consisted of 30 MRI sessions, each including a run of gradCPT performance, from a single individual (RTC, a 56-year-old left-handed male) collected over ten months. Across all sessions, the participant’s mean gradCPT *d*’ was 2.42 (range = 1.30–4.45; *s.d*. = .85). Performance increased monotonically over the course of the sessions (Spearman rank correlation between *d*’ and session number; *r_s_* = .53, *p* = .003). Due to hardware variability, mean gradCPT trial length ranged from 738 to 800 ms (mean = 778 ms, median = 794 ms, *s.d*. = 24.8 ms), and there was a non-significant monotonic relationship between trial length and *d*’, such that performance was numerically higher on gradCPT runs with a slower stimulus presentation rate (*r_s_* = .28, *p* = .14).

#### Trait-like attention prediction

We first validated the sustained attention CPM by applying it to predict this highly sampled individual’s average gradCPT performance, a trait-like measure of sustained attention abilities. When the model was applied to the participant’s mean gradCPT functional connectivity matrix across all 30 sessions, predicted *d*’ was remarkably similar to true average performance (predicted mean *d*’ = 2.37 vs. observed mean *d*’ = 2.42). Prediction error (i.e., the difference between observed and predicted mean *d*’ values) was smaller than the absolute prediction error of the task-based model for any participant in Rosenberg et al (2016a), despite the fact that the current data set is an external validation sample collected with different scan parameters (**Figure 5A**). The lower absolute error here may arise because the current data set includes more fMRI data per individual (165 min vs. 36 min) and/or reflects a more traitlike estimate of the participant’s overall ability to maintain focus. Furthermore, suggesting that prediction accuracy is not driven by regression to the training set mean, predicted mean *d*’ is closer to the participant’s true mean *d*’ than it is to the average *d*’ of all training subjects (|predicted mean *d*’ – observed mean *d*’| = .054; |predicted mean *d*’ – training set mean *d*’| = .26).

**Figure 5.**
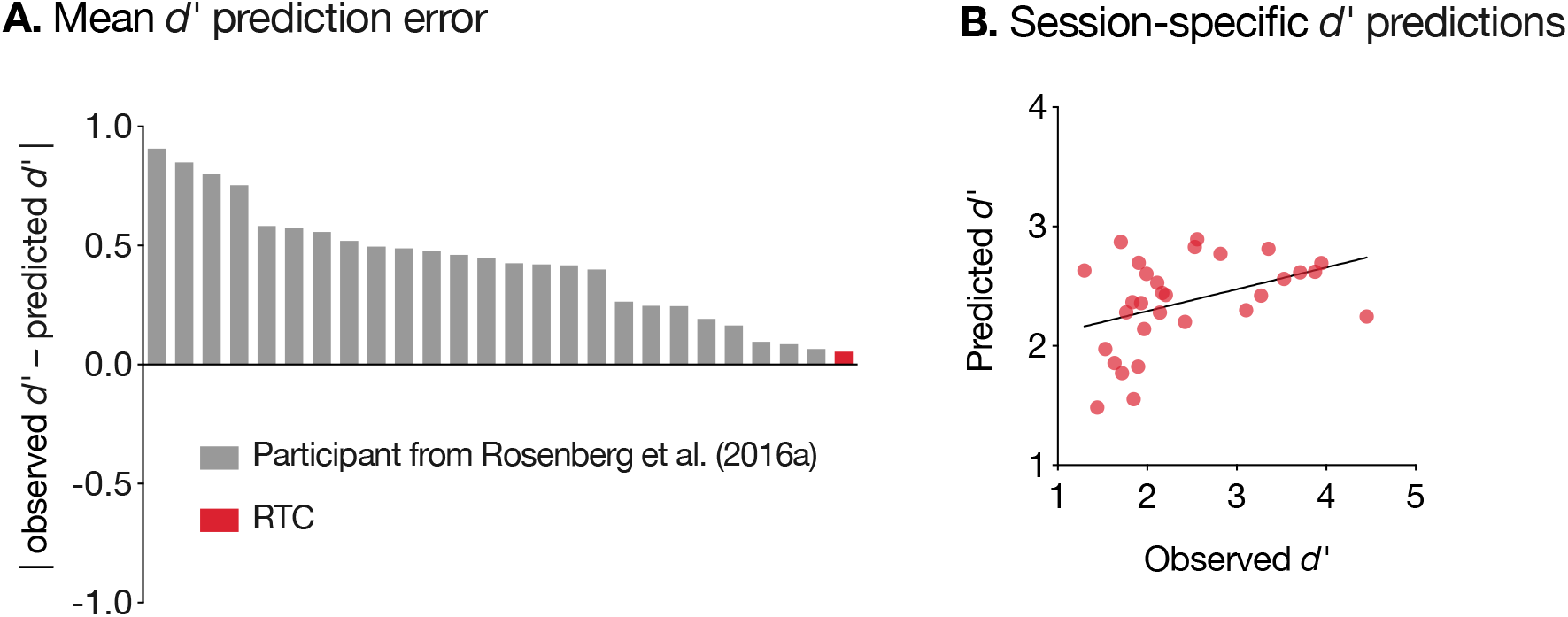
The sustained attention CPM generalized to predict an individual’s mean gradCPT performance (A) as well as day-to-day fluctuations around this mean (B).

#### State-like attention predictions

We next applied the sustained attention CPM to data from each scan session separately to generate session-specific *d*’ predictions. Predictions were positively correlated with observed *d*’ values (*r_s_* = .42, *p* = .02; **Figure 5B**), which reflect both state-like and trait-like aspects of sustained attention. Predictions remained significant when controlling for mean trial duration and session number with partial correlation (partial *r_s_* = .39, *p* = .04). As a post-hoc analysis, given that mean trial duration was non-normally distributed, we divided runs based on a median split of trial duration and assessed predictive power separately in each half. Predictions were significant in the 15 runs with faster trials (*r_s_* = .65, *p* = .0095) but non-significant in the 15 runs with slower trials (*r_s_* = .04, *p* = .88). Thus, the same networks that predict individual differences in attention, block-to-block fluctuations in an internal validation data set, and session-to-session variability in an independent 50-person sample capture mean performance and session-to-session fluctuations in a dense longitudinal phenotyping sample.

### Experiment 4: Propofol modulates sustained attention network strength

The sustained attention CPM captures within-subject changes in sustained attention across minutes, days, and weeks. These attentional state changes likely result from fluctuations in internal and external distraction, as well as variability in neurocognitive states such as motivation and sleepiness. To characterize the sensitivity of the sustained attention CPM to a completely different kind of attentional state change—one induced by pharmacological manipulation—we analyzed MRI data collected from 21 adults while awake (eyes-closed rest) and under deep sedation with propofol in the same imaging session (data set described in Qiu et al., 2017). End-tidal CO_2_, a measure of carbon dioxide concentration at the end of an exhalation, and heart rate did not significantly differ between the propofol and awake conditions. Mean blood pressure was lower in the propofol condition but fell within the autoregulatory range (Qiu et al., 2017).

As predicted, participants showed functional connectivity signatures of stronger attention—higher high-attention network strength and lower low-attention network strength— when awake than when under deep sedation (high-attention network: *t*_20_ = 4.53, *p* = 2.03×10^−4^ [effect larger than effects on 93.07% of same-size random networks]; low-attention network: *t*_20_ = –7.71, *p* = 2.05×10^−7^ [effect larger than effects on 99.98% of same-size random networks]; **Figure 6A**). In other words, the sustained attention CPM was sensitive to an anesthesia-induced attentional and cognitive state change.

**Figure 6.**
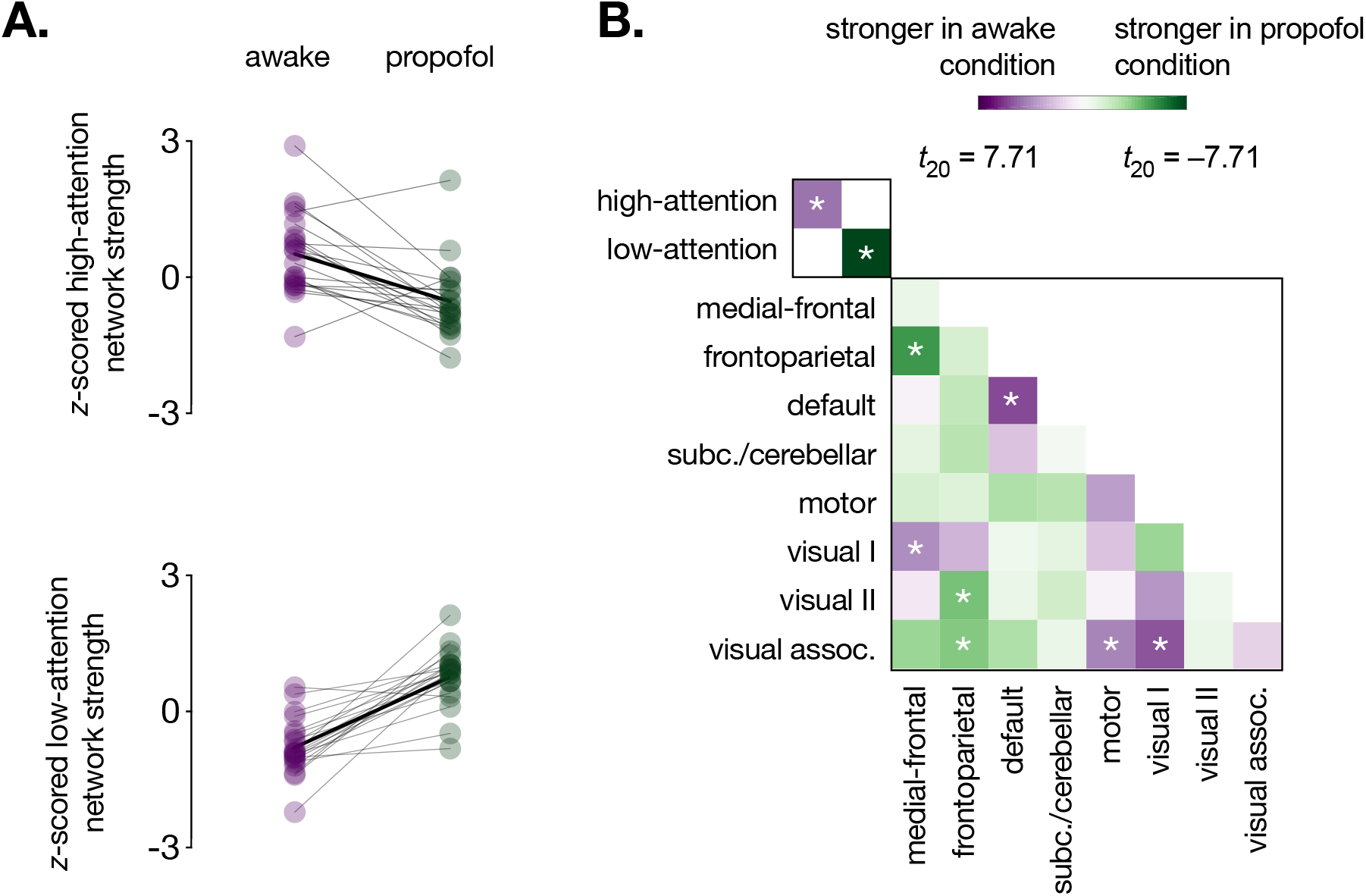
Effects of propofol on functional network strength. **A.** High-attention and low-attention network strength during the awake and deep sedation conditions. Network strength values were *z*-scored within graph for visualization. Individual dots represent individual participants, gray lines represent individual participant network strength change, and solid black lines indicate group mean change. **B.** Differences in within-network and between-network strength (i.e., summed functional connectivity) during the awake and deep sedation conditions. Low-attention network strength differed by condition more than any network pair in the lower right matrix; high-attention network strength differed more than 33/36 pairs. \**p* < .05/38

Demonstrating the relative specificity of propofol effects to *a priori* attention networks, the low-attention network showed a larger propofol effect than did any canonical resting-state network (defined in Finn et al., 2015). The high-attention network showed a larger propofol effect than all but within-default network connections, medial-frontal network-frontoparietal network connections, and visual I network-visual association network connections (**Figure 6B**).

### Experiment 5: Sevoflurane modulates sustained attention network strength

We replicated effects of anesthesia on the sustained attention CPM’s high- and low-attention networks using an independent sample of 11 adults scanned while awake, under light anesthesia with sevoflurane, and recovering from anesthesia (data set described in Martuzzi et al., 2010). Importantly, heart and respiratory rates, end-tidal CO_2_, and O_2_ partial pressure, a measure of blood oxygen saturation, did not significantly differ between the awake and sevoflurane conditions. Although systolic, diastolic, and mean blood pressure decreased under anesthesia, these changes fell within the autoregulatory range and were unlikely to have resulted in hemodynamic changes (Martuzzi et al., 2010).

As with propofol, participants showed higher high-attention and lower low-attention network strength in the awake than the anesthesia condition (high-attention network: *t*_10_ = 2.15, *p* = .057 [effect larger than effects on 85.15% of same-size random networks]; low-attention network: *t*_10_ = –3.37, *p* = .0071 [effect larger than effects on 96.32% of same-size random networks]; **Figure 7A**). High-attention network strength was higher during the awake than the recovery scan (*t*_10_ = 2.27, *p* = .047), and did not significantly differ between the anesthesia and the recovery scans (*t*_10_ = .57, *p* = .58). Low-attention network strength was not significantly different during the awake and recovery scans (*t*_10_ = 1.56, *p* = .15), but there was a trend such that it was lower during the recovery than the anesthesia scan (*t*_10_ = 2.14, *p* = .058). Wholebrain analyses revealed stronger connectivity between the medial-frontal and visual I networks, and the motor and visual association networks, during the awake than the anesthesia condition. Connectivity between the frontoparietal and visual association networks was significantly stronger during the anesthesia than the awake condition (**Figure 7B**).

**Figure 7.**
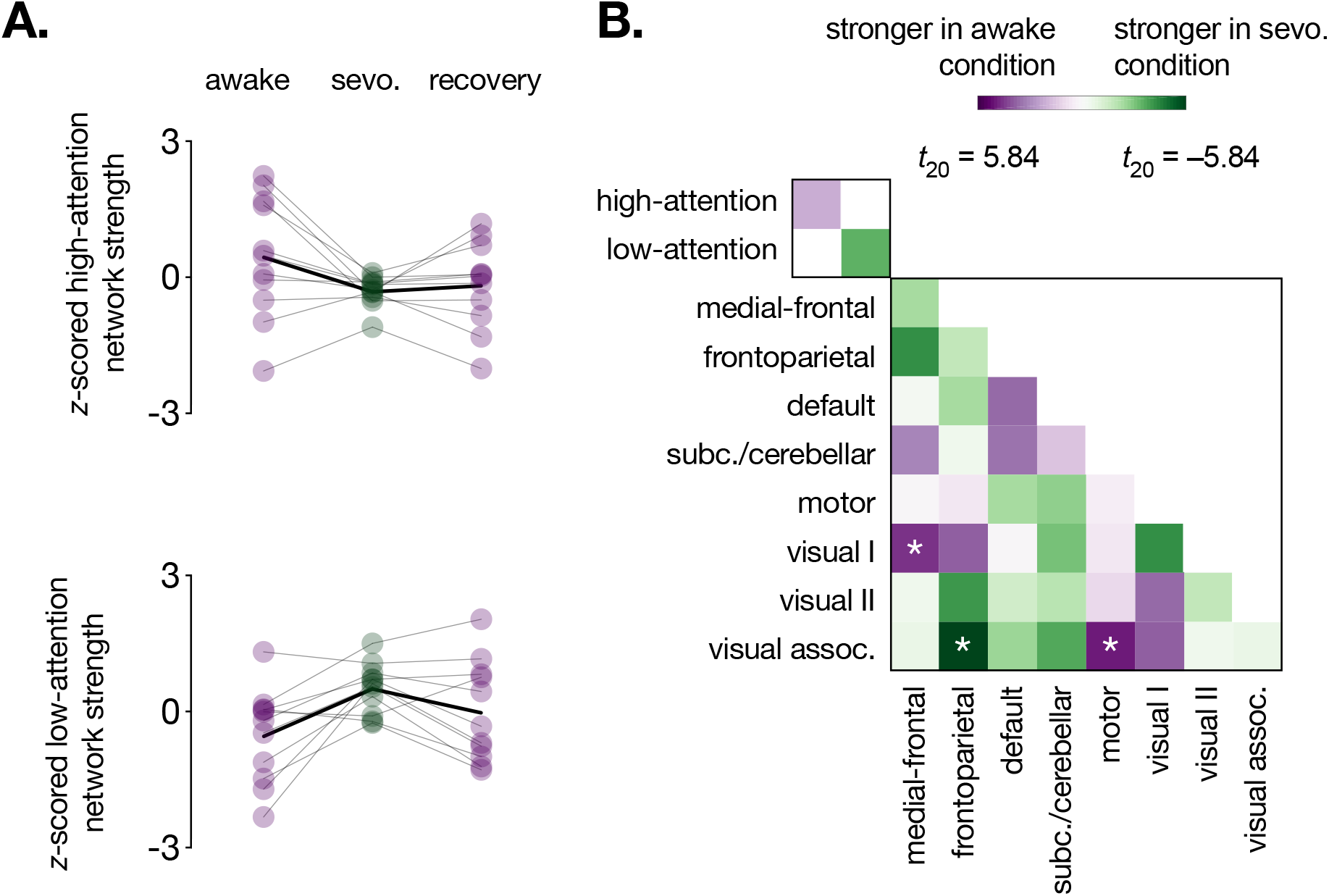
Effects of sevoflurane on functional network strength. **A.** High-attention and low-attention network strength during the awake (pre-anesthesia), sevoflurane, and recovery (postanesthesia) conditions. Network strength values were *z*-scored within graph for visualization. Individual dots represent individual participants, gray lines represent individual participant network strength change, and solid black lines indicate group mean change. **B.** Differences in within-network and between-network strength (i.e., summed functional connectivity) during the awake (pre-anesthesia) and sevoflurane conditions. \**p* < .05/38

#### Network strength as a function of state change

Propofol and sevoflurane are different anesthetic agents with different pharmacodynamic effects. Whereas participants in the propofol study did not respond to verbal call during the anesthesia condition (Experiment 4), participants in the sevoflurane study were only under light anesthesia (Experiment 5). To characterize relationships between the degree of anesthesia-induced cognitive and attentional state change and attention network strength, we generated a pseudo-dose-response curve by collapsing data across studies. State affected high-attention (*b* = –.57, *SE* = .15, *F*(3,48.9) = 9.61, *p* = 4.29×10^−5^) and low-attention (*b* = .91, *SE* = .14, *F*(3,2.0) = 24.64, *p* = .039) network strength, such that high-attention network strength systematically decreased and low-attention network strength systematically increased as state changes became more dramatic (**Figure 8**).

**Figure 8.**
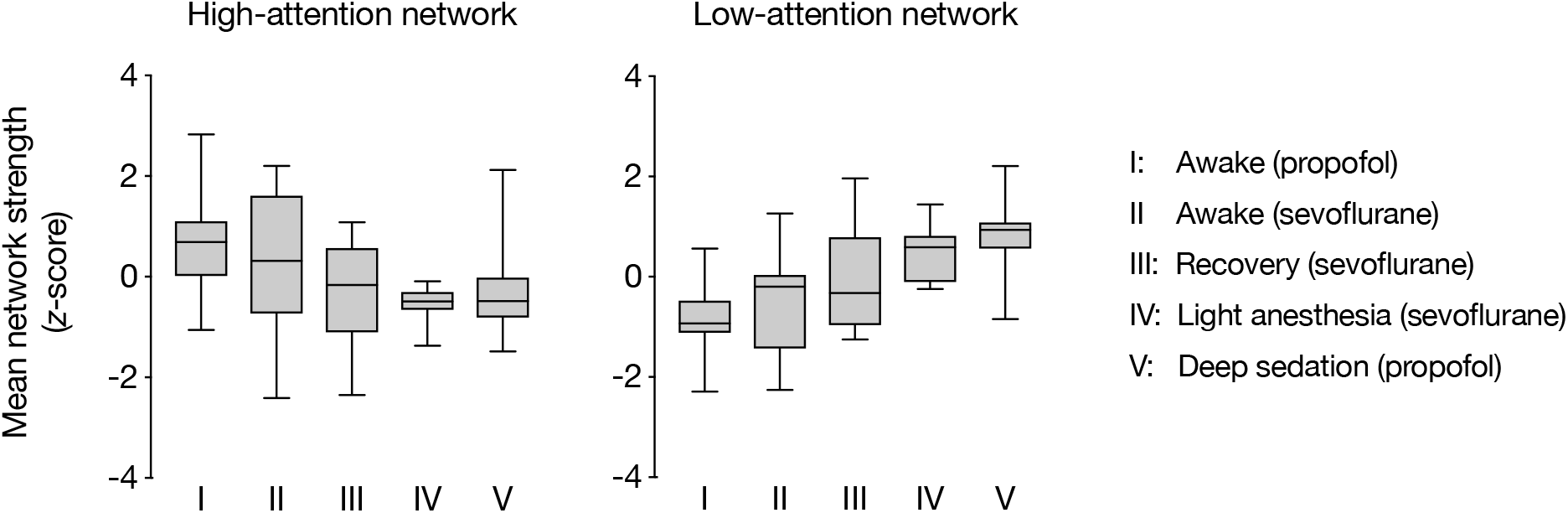
“Dose-response” curve relating attention network strength to the intensity of state changes across two data sets. Raw network strength values (summed Fisher *z*-transformed correlation coefficients) were normalized within the high-attention network and low-attention network plots separately. Boxes extend from the 25^th^ to 75^th^ percentiles, and whiskers from the minimum to maximum values. Horizontal lines correspond to group medians. State changes were most pronounced in the “Deep sedation (propofol)” condition, in which participants were under deep sedation and did not respond to verbal call. Effects of anesthesia were less pronounced in the “Light anesthesia (sevoflurane)” condition, and lesser still in the “Recovery (sevoflurane)” condition. Pre-anesthesia conditions, “Awake (propofol)” and “Awake (sevoflurane)”, are equivalent to task-free resting-state scans.

## Discussion

Each person has a unique pattern of functional brain connectivity, that, like a fingerprint, distinguishes them from a group (Finn et al., 2015; Miranda-Dominguez et al., 2014). Unlike a fingerprint, however, this pattern predicts cognitive and attentional abilities (Dubois and Adolphs, 2016; Rosenberg et al., 2017) and changes on multiple time scales (i.e., minutes, hours, days, development). Are these changes meaningful—that is, do they reflect behaviorally relevant changes in cognitive and attentional states?

Here we tested whether a publication-preregistered model, the sustained attention connectome-based predictive model, predicts both trait-like and state-like measures of attention. An internal validation (i.e., leave-one-subject-out) approach first revealed that functional connectivity patterns observed during 3-minute task blocks and 32-second rest breaks predict individuals’ block-specific task performance (Experiment 1). In other words, the same models that predicted participants’ average task performance in previous work (Rosenberg et al., 2016a) were also sensitive to block-to-block fluctuations in performance. When applied to a 50-person external validation sample, a model defined using the full Experiment 1 data set—the sustained attention CPM—not only predicted participants’ task performance during two fMRI sessions, but also predicted which session had better performance and which session had worse (Experiment 2). Moreover, when the sustained attention CPM was applied to data from a one individual’s 30 fMRI sessions, the prediction based on his average functional connectivity pattern reflected his average task performance, and predictions based on session-specific patterns reflected session-specific task performance (Experiment 3). Finally, the sustained attention CPM’s high- and low-attention networks were modulated by sevoflurane and propofol, such that functional connectivity signatures of better attention were observed when individuals were awake than when they were under light anesthesia or deep sedation (Experiments 4 and 5). Together these findings demonstrate that behaviorally relevant attentional states are reflected in functional connectivity patterns calculated from less than 30 seconds of rest data; that, when averaged over many scan sessions, functional connectivity patterns provide near-perfect predictions of average sustained attention function; and that the same functional connections that vary with sustained attention *across* individuals also change with attentional state (pharmacologically induced or not) *within* individuals.

This work aligns with a growing body of evidence that changes in ongoing attention and cognition are reflected in changes in functional connectivity. In particular, functional connectivity dynamics as measured by fMRI—particularly in the default mode and dorsal attention networks—mirror changes in task state (Gonzalez-Castillo et al., 2015; Shirer et al., 2011), task performance (Kucyi et al., 2016b; Shappell et al., 2019; Shine et al., 2016a, 2016b), and selfreported mind wandering (Kucyi, 2017; Kucyi and Davis, 2014; Turnbull et al., 2019). Furthermore, a recent intracranial electroencephalography study in humans revealed that default and dorsal attention network activity was more anticorrelated during periods of better attention task performance, and that dorsal attention activations preceded (and potentially caused) default mode deactivations (Kucyi et al., 2018). Complementing these findings, our results demonstrate for the first time that the same functional networks that *predict* individual differences in attention in novel individuals also predict attentional states specific to task blocks, fMRI sessions, and states of consciousness induced by anesthesia. Furthermore, they suggest that attentional-state-relevant dynamics are not constrained to an *a priori* set of canonical functional networks (e.g., the default mode, dorsal attention, and salience networks, which do not dominate the sustained attention CPM’s high- and low-attention networks; Rosenberg et al., 2016a), but rather span a distributed set of cortical, subcortical, and cerebellar brain regions (**Figure 3**).

At the same time as evidence for behaviorally meaningful functional connectivity dynamics accumulates, work shows that functional connection reliability is poor when measured in short time windows (even up to 36 minutes of data from a single fMRI session; Noble et al., 2017) and that dynamics arise due to sources including motion and sampling variability (Laumann et al., 2016). How do we resolve these discrepant observations? One possibility is that attentional-state-specific functional connectivity patterns have been previously characterized as noise. That is, attention changes over multiple time scales, and these changes are likely missed by analyses that group task states regardless of behavioral performance or average performance over long periods of time. A corollary to this suggestion is that functional connectivity patterns averaged over long periods of time and multiple fMRI sessions better approximate a person’s “true” connectome and better predict behavior in part because they sample a wider range of a person’s possible cognitive and attentional states. Importantly, we are not discounting the serious and well-documented effects of physiological and measurement noise on functional connectivity measured at short time scales. Rather, we suggest that a person’s attentional state is another statistically significant source of variability in data-driven functional networks that predict attention. (Simultaneously, networks that predict other processes such as emotional reactivity may fluctuate with changes in these states as well.) In the future, multi-session, multi-task fMRI samples and high-frequency behavioral sampling (e.g., ongoing task performance, pupillometry) can help disentangle the contributions of multiple sources of functional connectivity dynamics.

At first glance, the finding that functional network connectivity predicts moment-to-moment and day-to-day differences in attention stands in apparent contrast to work showing that the organization of canonical functional brain networks reflects stable individual differences rather than task states or day-to-day variability (Gratton et al., 2018). However, the results are not incongruous. First, Gratton et al. characterized the topography of canonical networks such as the default and dorsal attention networks, whereas we consider the strength of a distributed set of functional connections selected to predict behavior with a data-driven approach. More importantly, however, Gratton and colleagues found that, while task states and fMRI sessions are not dominant sources of variability in functional network organization, they do have significant effects. Thus, it is possible that collapsing across attentional states within broader task states obscures behaviorally relevant differences in network organization, and that characterizing networks during distinct periods of successful and unsuccessful task performance could magnify these small-but-significant effects. Future work characterizing the effects of task challenges and pharmacological interventions (extrinsic state manipulations) as well as attention fluctuations (intrinsic state manipulations) on functional network organization and connectivity patterns will further inform the sources of variability around each individual’s average functional connectome fingerprint.

Together the current work demonstrates that transient functional connectivity patterns reflect local attentional states. However, it remains an open question whether task-based or resting-state functional connectivity patterns predict changes in attention over longer periods of time, such as development and aging. Looking ahead, testing the sensitivity of connectome-based models to within-subject changes in attentional and cognitive abilities over years and decades can inform the common and distinct functional architecture of these processes across the lifespan (Rosenberg et al., 2018b). Furthermore, building new models to predict developmental trajectories in abilities and behavior can provide insights into the ways in which functional brain organization reflects risk for or resilience to impairments such as attention deficit hyperactivity disorder, potentially informing early treatments or interventions.

In sum, we show that a neuromarker of sustained attention generalizes across five independent data sets to predict individual differences in sustained attention as well as intrinsic and pharmacologically induced attentional states from task-based and task-free functional connectivity. Thus, functional connectivity patterns reflect a combination of trait-like and state-like aspects of attention, and, more broadly, dynamics in functional connectivity in part reflect dynamics in attentional and cognitive states.

## Acknowledgments

This work was supported by the Yale FAS MRI Program funded by the Office of the Provost and the Department of Psychology; a National Science Foundation Graduate Research Fellowship, American Psychological Association Dissertation Research Award, and Theresa Seessel Postdoctoral Fellowship to M.D.R.; NIH Medical Scientist Training Program Training Grant T32GM00720 supporting A.S.G.; National Science Foundation BCS1558497 to M.M.C., and National Institutes of Health MH108591 to M.M.C., EB009666 to R.T.C., and T32 DA022975 to D.S.

## Author Contributions

Conceptualization, M.D.R., M.M.C., D.S., E.S.F., and R.T.C.; Methodology, M.D.R., E.S.F., and D.S.; Formal Analysis, M.D.R; Investigation, M.D.R., D.S., A.S.G., E.W.A., Y.H.K, R.R., and M.Q.; Writing – Original Draft, M.D.R.; Writing – Review & Editing, M.M.C., D.S., A.S.G., E.W.A., Y.H.K., E.S.F., R.R., M.Q., and R.T.C.; Funding Acquisition, M.M.C. and R.T.C.; Resources, M.D.R, D.S., R.T.C, and M.M.C.; Supervision, M.M.C., D.S., and R.T.C.

## Methods

### Experiment 1: Sustained attention network strength predicts minute-to-minute attention fluctuations

#### Participants

Thirty-one right-handed, neurologically healthy adults with normal or corrected-to-normal vision were recruited from Yale University and the surrounding community to perform the gradual-onset continuous performance task (gradCPT; Esterman et al., 2014; Rosenberg et al., 2013) and rest during MRI data collection (data set described in detail in Rosenberg et al., 2016). All provided written informed consent and were paid for their participation. Following exclusion for excessive head motion (greater than 2 mm translation or 3° rotation in all functional runs) or insufficient coverage, data from 25 individuals were submitted to further analysis (13 females, 18–32 years, mean = 22.7 years).

#### Experimental design

The gradCPT, a test of sustained attention and inhibitory control, was used to assess participants’ overall ability to maintain focus and to track their attention fluctuations over time. During the gradCPT, grayscale images of city (90%) and mountain (10%) scenes gradually transitioned from one to the next every 800 ms. Participants were instructed to press a button with their right index finger every time they saw a city scene, but not to press to mountain images. Because stimuli were constantly in transition, an iterative algorithm was used to assign button press responses to trials (Rosenberg et al., 2016a).

Scan sessions began with a high-resolution anatomical image acquisition followed by a 6-min resting-state run, three 13:44-min gradCPT runs, and a second 6-min resting-state run. GradCPT runs included 8-s of fixation followed by four 3-min task blocks interleaved with 32-s rest breaks. Resting-state runs are not analyzed here.

For each task block (3 runs x 4 blocks/run = 12 blocks total), performance was measured with sensitivity (*d*’), or the inverse of the standard normal cumulative distribution function of the false alarm rate (incorrect presses to mountains) subtracted from the inverse of the standard normal cumulative distribution function of the hit rate (correct presses to cities). Overall gradCPT performance was measures by averaging *d*’ values across blocks.

#### Imaging parameters and preprocessing

MRI data were collected at the Yale Magnetic Resonance Research Center on a 3T Siemens Trio TIM system using a 32-channel head coil. Functional runs included 824 (task) or 363 (rest) whole-brain volumes acquired using a multiband echo-planar imaging (EPI) sequence with the following parameters: repetition time (TR) = 1,000 ms, echo time (TE) = 30 ms, flip angle = 62°, acquisition matrix = 84 × 84, in-plane resolution = 2.5 mm^2^, 51 axial-oblique slices parallel to the AC-PC line, slice thickness = 2.5 mm, multiband 3, acceleration factor = 2. Parameters of the anatomical magnetization prepared rapid gradient echo (MPRAGE) sequence were as follows: TR = 2530 ms, TE = 3.32 ms, flip angle = 7°, acquisition matrix = 256 × 256, in-plane resolution = 1.0 mm^2^, slice thickness = 1.0 mm, 176 sagittal slices. A 2D T1-weighted image coplanar to the functional images was also collected for registration.

Functional data were analyzed using BioImage Suite (Joshi et al., 2011) and custom Matlab scripts (Mathworks) as described previously (Rosenberg et al., 2016a). Motion correction was performed using SPM8. Linear and quadratic drift, mean signal from the cerebrospinal fluid (CSF) and white matter, and 24 motion parameters were regressed from the data. Global signal was also included as a nuisance regressor to reduce the confounding effects of motion (Ciric et al., 2017; Rosenberg et al., 2018a). Data were temporally smoothed with a zero mean unit variance Gaussian filter (cutoff frequency = 0.12 Hz).

Preprocessing steps were applied to data concatenated across task runs as well as to data from each task block and rest break separately to maintain independence between block- and break-specific functional connectivity matrices (see the *Functional connectivity matrix generation* section for detail).

Because head motion can confound functional connectivity analyses, five task runs and two rest runs with more than 2 mm head translation or 3° head rotation were excluded from further analysis. As reported previously, mean frame-to-frame head displacement during gradCPT runs and average motion across gradCPT runs did not correlate with *d*’ across individuals (|*r*| values ≤ .1; *p* values ≥ .62; Rosenberg et al., 2016a).

#### Functional connectivity matrix generation

Functional network nodes were defined with a 268-node atlas that includes cortical, subcortical, and cerebellar nodes (Shen et al., 2013). The atlas was warped from MNI space into single-subject space via linear and nonlinear registrations between the EPI images, coplanar scan, 3D anatomical scan, and MNI brain.

To generate whole-brain functional connectivity matrices, the mean fMRI signal timecourse for each node was calculated by averaging the timecourses of its constituent voxels. The Pearson correlation between the average timecourses of every pair of nodes was computed and Fisher *z*-transformed to yield symmetrical 268 × 268 matrices of functional connections, or edges.

To test whether the same functional networks that predict individual differences in attention also predict attention fluctuations, we measured participants’ *overall* pattern of functional connectivity during task engagement as well as their connectivity patterns during shorter intervals over the course of the task. To this end, for each participant we calculated *(A)* a single overall task matrix from data concatenated across task runs, excluding volumes collected during rest breaks; *(B)* up to 12 task-block matrices from volumes acquired during individual task blocks; and *(C)* up to 9 rest-break matrices using volumes acquired during individual rest breaks (starting 6 s after break onset and ending 3 s before the onset of the upcoming task block to reduce the influence of task-related stimulus processing on rest-break connectivity).

#### Connectome-based predictive modeling

Previous analyses of these data demonstrated that models based on patterns of functional connectivity observed during the gradCPT—here, the “overall task matrices”—predict individual differences in performance from both task-based and resting-state functional connectivity (Rosenberg et al., 2016a). Furthermore, these same network models generalize to independent data sets to predict measures of attention and inhibitory control including attention deficit hyperactivity disorder symptoms, stop-signal task performance, and Attention Network Task performance (Rosenberg et al., 2016a, 2016b; Rosenberg et al., 2018).

To test whether these same models predict not only differences in attention between individuals but also differences in attention within single individuals over time, we performed a variant of connectome-based predictive modeling (Shen et al., 2017) using leave-one-subject-out cross-validation. In this pipeline, feature selection and model building steps replicate those described in Rosenberg et al. (2016a), but models are tested on different data from the held-out individual.

First, data from one individual were set aside, leaving overall task matrices and overall *d*’ scores from the remaining 24 participants. Next, robust regression between each edge in the overall task matrices (35,778 total) and *d*’ was performed across subjects. Edges related to behavior at *p* < .01 were retained and separated into a positive tail (positive regression coefficients) and a negative tail (negative regression coefficients).

For each participant in the training set, overall strength in the positive and negative tails was calculated by summing their respective connections. The difference in connectivity strength between the tails (positive tail strength – negative tail strength) was used as a predictor in a linear regression of the form *d*’ = a*X* + b. This model differs slightly from the general linear model reported in Rosenberg et al. 2016a, in which positive and negative tail strength were included as independent predictors. Here the difference in strength between the tails is used to avoid collinear predictors.

To test whether this model generalized to predict attention fluctuations in previously unseen individuals, it was applied to each of the left-out participant’s task-block matrices separately. In other words, the difference between positive and negative tail strength was calculated using data from each of the held-out individual’s 3-min task blocks. The resulting difference scores (8 for participants with two gradCPT runs and 12 for participants with three) were input in the model to generate a predicted *d*’ value for each task block. Model performance was assessed by computing the Spearman correlation between the left-out individual’s predicted and observed block-specific *d*’ scores.

If changes in task-based functional connectivity predict changes in attention within subjects, task-free connectivity patterns may also reflect transient attentional states. To test this possibility, the same model was applied to each of the left-out subject’s rest-break matrices to generate a predicted *d*’ score corresponding to each rest break (6 for participants with two gradCPT runs and 9 for participants with three). Model performance was assessed by correlating predictions with the *d*’ scores from the preceding and following task blocks since performance is not measured during the breaks themselves. These models are referred to as the “pre-break” and “post-break” models, respectively, throughout the text.

Finally, the prediction pipeline was repeated until every individual had been left out once.

#### Significance testing

The significance of model predictions was evaluated at both the individual and group levels. To determine whether models significantly predicted fluctuating *d*’ in a single person, block-specific *d*’ scores were shuffled 1,000 times and correlated with model predictions. This process generated two null *r_s_*-value distributions per individual: one of null task-block predictions and one of null rest-break predictions. The significance of task-block model predictions was calculated as *p* = (1+ the number of null task-block *r_s_*-values ≥ the observed task-block *r_s_*-value)/1001. The significance of rest-break model predictions was calculated as *p* = (1+ the number of null rest-break values ≥ the observed pre-break *r_s_*-value)/1001 and *p* = (1+ the number of null rest-break values ≥ the observed post-break *r_s_*-value)/1001.

The significance of model predictions was also evaluated at the group-level with paired *t*-tests comparing observed within-subject *r_s_*-values to the mean of each participant’s null task-based or rest-break *r_s_*-value distribution. Spearman correlation coefficients were Fisher *z*-transformed before averaging; averaged *z*-values were converted back to *r_s_*-values for reporting.

### Experiment 2: Sustained attention network strength predicts session-to-session changes in attention task performance

#### Participants and experimental design

98 individuals (62 female, age 18–35, mean = 22.9 years, *s.d*. = 4.6 years) participated in a two-session neuroimaging experiment designed to assess different aspects of attention and memory. 90 participants completed both sessions, which included 10-minute resting-state runs (two in session 1 and one in session 2); an Inscapes movie run (one per session; Vanderwal et al., 2015); gradCPT, multiple object tracking, visual short-term memory task runs (one per session; order counterbalanced across participants and sessions); and an Attention Network Task run (one in session 2). All participants provided written informed consent in compliance with procedures approved by the Yale University Human Subjects Committee and were paid for their participation.

GradCPT data from the 50 individuals with whole-brain coverage, *d*’ scores within 3 standard deviations of the group mean, and acceptable levels of head motion (<3 mm maximum head displacement and <.15 mm mean frame-to-frame displacement) during session 1 and session 2 were analyzed here (34 female, age 18–32, mean = 23.3 years, *s.d*. = 4.1 years). For these individuals, fMRI sessions were held approximately three weeks apart (range = 5–133 days, mean = 20.2 days, *s.d*. = 24.9 days). The number of days separating sessions 1 and 2 did not correlate with gradCPT performance on either day or with the difference between them (|*r_s_*| values < .0482, *p* values > .73). No other data were tested, and this data set has not been published previously.

#### Imaging parameters and preprocessing

MRI data were collected at the Yale Magnetic Resonance Research Center on a 3T Siemens Prisma system using a 64-channel head coil. Functional gradCPT runs included 600 whole-brain volumes acquired using a multiband EPI sequence with the following parameters: TR = 1,000 ms, TE = 30 ms, flip angle = 62°, acquisition matrix = 84 × 84, in-plane resolution = 2.5 mm^2^, 52 axial-oblique slices parallel to the AC-PC line, slice thickness = 2.5 mm, multiband 4, acceleration factor = 1. Parameters of the MPRAGE sequence were as follows: TR = 1800 ms, TE = 2.26 ms, flip angle = 8°, acquisition matrix = 256 × 256, in-plane resolution = 1.0 mm^2^, slice thickness = 1.0 mm, 208 sagittal slices.

Data were processed with AFNI (Cox, 1996). Preprocessing steps included the exclusion of three volumes from the start of each run; censoring of volumes in which more than 10% of voxels were outliers; censoring of volumes for which the Euclidean norm of the head motion parameter derivatives exceeded .2; despiking; slice-time correction; motion correction; and regression of mean signal from the CSF, white matter, and whole brain as well as 24 motion parameters. Functional images were aligned to their corresponding skull-stripped high-resolution anatomical image via linear transformation, and the anatomical image was aligned to MNI space. For each session, a functional connectivity matrix was defined from gradCPT data as described in Experiment 1.

Because head motion and amount of data available after censoring can confound functional connectivity analyses, we confirmed that *d*’ values were not significantly correlated with mean frame-to-frame motion after censoring, maximum motion after censoring, or number of frames censored (session 1: |*r*| values < .15, *p* values > .31; session 2: |*r*| values < .17, *p* values > .26). Head motion and number of post-censoring volumes did not differ between participants’ day 1 and day 2 gradCPT runs, their better and worse gradCPT runs, or their predicted-better and predicted-worse gradCPT runs (|*t*_49_| statistics < 1.24, *p* values > .22).

#### Attention predictions

The model used for external validation, the sustained attention connectome-based predictive model (Rosenberg et al., 2016a), was defined using the high-attention and low-attention networks described in the *Predictive network anatomy* section of Experiment 1. Using data from all 25 participants in the Rosenberg et al. (2016a) sample, a linear regression of the form *d*’ = a*X* + b was computed where *X* = high-attention network strength – low attention network strength. This model, the sustained attention CPM, was applied, completely unchanged, to each participant’s session 1 and session 2 gradCPT matrix to generate session-specific *d*’ predictions. Model performance was assessed by rank correlating predictions with *d*’ scores across individuals for each session separately. A paired *t*-test was applied to compare predictions for participants’ better vs. worse session.

### Experiment 3: Sustained attention network strength predicts week-to-week changes in attention in a single individual

#### Participant and experimental design

30 sessions of MRI data were collected over ten months from a single individual (RTC, a 56-year-old left-handed male; Salehi et al., 2018). Data were acquired at the Yale Magnetic Resonance Research Center on two identically configured Siemens 3T Prisma scanners using a 64-channel head coil.

During each scan session, six 6:49-min task runs and two 6:49-min resting-state functional MRI runs (including initial shim time and 8 s of discarded acquisitions) were collected. Tasks included the gradCPT (450 trials/run) as well as an *n*-back task, stop-signal task, card guessing task, “reading the mind in the eyes” task, and movies task (described in detail in Salehi et al., 2018). Here we restrict our analyses to gradCPT data. Attention was operationalized as gradCPT sensitivity (*d*’).

#### Imaging parameters and preprocessing

A high-resolution anatomical (MPRAGE) scan was collected during the first session with the following parameters: 208 contiguous slices acquired in the sagittal plane, TR = 2400 ms, TE = 1.22 ms, flip angle = 8°, slice thickness = 1 mm, inplane resolution = 1 mm^2^, matrix size = 256×256. Functional images were collected using a multiband gradient EPI sequence with the following parameters: 75 contiguous slices acquired in the axial-oblique plane parallel to AC-PC line, TR = 1000 ms, TE = 30 ms, flip angle = 55°, slice thickness = 2 mm, multiband = 5, acceleration factor = 2, in-plane resolution = 2 mm^2^, matrix size = 110×110 (Salehi et al., 2018).

Imaging data were analyzed using BioImage Suite and custom MATLAB scripts. Motion correction was performed using SPM12. White matter and CSF masks were defined in MNI space and warped into single-subject space using linear and nonlinear transformations. Linear, quadratic, and cubic drift, a 24-parameter motion model, mean signal from CSF and white matter, and mean global signal were regressed from the data. Lastly data were temporally smoothed with a Gaussian filter (sigma = 1.55, cutoff frequency = 0.121 Hz; see Salehi et al., 2018 for detail).

For each session, a functional connectivity matrix was generated from gradCPT data as described in Experiment 1. Due to hardware variability, mean gradCPT trial length ranged from 738 to 800 ms (mean = 778 ms, median = 794 ms, *s.d*. = 24.8 ms). Thus, matrices were calculated from the first 330 volumes of every run to account for differences in task duration.

#### Attention predictions

The sustained attention CPM was applied to the participant’s gradCPT functional connectivity matrices to generate session-specific *d*’ predictions. Model performance was assessed by rank correlating predictions with *d*’ scores across the 30 sessions.

### Experiment 4: Propofol alters sustained attention network strength

#### Participants and experimental design

Functional MRI data were collected from 32 adults while awake (eyes-closed rest) and under deep sedation in accordance with research protocols approved by the Yale University Institutional Review Board (age 19–35 years, 13 female; data set described in Qiu et al., 2017). After exclusion for excessive head motion (>.15 mm mean frame-to-frame displacement) in either condition, data from 21 participants remained. Head motion did not differ between awake and deep sedation conditions (awake: 0.077 ± 0.04 mm, deep sedation: 0.077 ± 0.03 mm, *p* = 0.98; Qiu et al., 2017).

#### Imaging parameters and preprocessing

MRI data were collected at the Yale Magnetic Resonance Research Center on a 3T Siemens Trio TIM system using a 12-channel head coil. Two functional runs collected during both the awake and deep sedation conditions included 210 whole-brain volumes acquired using an EPI sequence with the following parameters: TR = 2000 ms, TE = 30 ms, field-of-view (FOV) = 256×256 mm^2^, flip angle = 90°, matrix size = 64×64, 33 AC-PC aligned slices, slice thickness = 4 mm, no gap. After data were collected during the awake condition, propofol was infused intravenously to induce an anesthetized state (“deep sedation”) in which participants did not respond to verbal call. During propofol infusion, a high-resolution MPRAGE scan was collected with the following parameters: 176 contiguous sagittal slices, voxel size = 1 mm^3^, FOV = 256×256 mm^2^, TR = 2530 ms, TE = 3.34 ms, flip angle = 7°. A 2D T1-weighted image with the following parameters was also acquired for the purpose of registration: TR = 300 ms, TE = 2.43 ms, FOV = 256×256 mm^2^, matrix size = 256 × 256, flip angle = 60°.

Imaging data were analyzed with SPM8 (slice-time correction and motion correction), BioImage Suite (all other preprocessing steps), and custom Matlab scripts. After the first ten volumes of each functional run were discarded, data were slice-time corrected, motion corrected, and iteratively smoothed to a smoothness of approximately 6 mm full width half maximum. Covariates of no interest were regressed from the data including linear and quadratic drift, mean CSF, WM, and gray matter signal, and a 24-parameter motion model. Data were temporally smoothed with a Gaussian filter (cutoff frequency = approximately 0.12 Hz). For additional detail see Qiu et al. (2017). Time-series data were concatenated across the two runs from the same condition. Functional connectivity matrices were calculated from the awake and propofol conditions as described in Experiment 1.

#### Network strength comparison

A paired *t*-test was used to compare high-attention and low-attention network strength in the awake and deep sedation conditions. Differences in high-attention and low-attention network strength were compared to differences in 10,000 same-size random networks. To further assess the specificity of state-dependent differences, we also compared propofol effects on high-attention and low-attention network strength to propofol effects on functional connectivity within and between canonical resting-state networks including the medial-frontal, frontoparietal, default mode, subcortical-cerebellar, motor, visual I, visual II, and visual association networks (Finn et al., 2015).

### Experiment 5: Sevoflurane alters sustained attention network strength

#### Participants and experimental design

Neuroimaging data were collected in a single imaging session from 14 adults (7 female, age 22–34) while awake, under light anesthesia, and during recovery after anesthesia (data set described in Martuzzi et al., 2010). Data from three participants were excluded from analysis due to excessive head motion during scanning (>0.15 mm frame-to-frame displacement). During the pre-anesthesia awake condition, participants received pure oxygen through a facemask. During the sevoflurane condition, participants received a mixture of oxygen and sevoflurane (end-tidal concentration 1%, equivalent to 0.5 minimum alveolar concentration). In the post-anesthesia recovery condition, participants received pure oxygen again. Conditions were separated by 10-min transition periods, and participants were instructed to keep their eyes closed throughout the study. All participants gave written informed consent and the Human Investigation Committee of the Yale School of Medicine approved the study protocol.

#### Imaging parameters and preprocessing

MRI data were collected at the Yale Magnetic Resonance Research Center on a 3T Siemens Trio TIM system using a circularly polarized head coil (Martuzzi et al., 2010). Scan sessions began with a localizer followed by a 2D T1-weighted anatomical scan (TR = 300 ms, TE = 2.43 ms, FOV = 256 mm, matrix size = 256 × 256, flip angle 60°, 33 axial slices parallel to the AC-PC line, slice thickness = 4 mm, no gap). Functional runs included 210 volumes and were collected during the pre-anesthesia, anesthesia, and post-anesthesia conditions using a T2_*_-sensitive gradient-recalled, single-shot echo-planar imaging pulse sequence (TR = 2 s, TE = 31 ms, FOV = 256 mm, flip angle 90°, matrix size 64 × 64, 33 slices parallel to the bicommissural plane, slice thickness = 4 mm, no gap). High-resolution anatomical (MPRAGE) images were acquired in between the anesthesia and post-anesthesia conditions (176 contiguous sagittal slices, slice thickness = 1 mm, matrix size 256 × 256, FOV = 256 mm TR = 2530 ms, TE = 3.34 ms, flip angle = 7°).

Data were analyzed with BioImage Suite. After the first ten volumes of each functional run were discarded, data were temporally and spatially realigned, corrected to remove slice means and drift, and low-pass filtered at a cut-off frequency of 0.08 Hz. Covariates of no interest were regressed from the data including mean CSF and white matter signal and a 6-parameter motion model. Functional images were co-registered to the 2D anatomical image. The 2D anatomical image was then registered to the 3D anatomical image, and the 3D anatomical image was aligned into MNI reference space via non-linear transformation (Martuzzi et al., 2010). Functional connectivity matrices were calculated from the awake, sevoflurane, and recovery conditions as described in Experiment 1.

#### Network strength comparison

Changes in high- and low-attention network strength as well as canonical resting-state networks were assessed as described in Experiment 4.

#### Network strength as a function of state change

The relationship between cognitive state and normalized attention network strength was assessed with a linear mixed effects model using the lme4 package (Bates et al., 2015) in R. State, an ordered factor with levels {rest, recovery, sevoflurane, propofol}, was entered into the model as a fixed effect. Intercepts for data sets and participants nested within data set were included as random effects. The limited-memory Broyden–Fletcher–Goldfarb–Shanno algorithm (L-BFGS-B) (Byrd et al., 1995), implemented with the optimx package (Nash and Varadhan, 2011), was used for optimization. *P*-values were obtained using Type III Satterthwaite approximations with the lmerTest package (Kuznetsova et al., 2017).

